# Angle-tuned Coil: A Focal Spot-size Adjustable Transcranial Magnetic Stimulator

**DOI:** 10.1101/771758

**Authors:** Qinglei Meng, Hedyeh Bagherzadeh, Julian Loiacono, Xiaoming Du, Elliot Hong, Yihong Yang, Hanbing Lu, Fow-Sen Choa

## Abstract

**Background:** Conventional transcranial magnetic stimulation (TMS) coils are limited by the depth-focality tradeoff rule and the emission field intensity from coils with either small or large apertures will diverge quickly at less than one aperture diameter distance away from the coil. To utilize a better depth-focality tradeoff rule and accomplish deep and focused stimulation, a new approach needs to be employed.

**Objectives:** We report a new TMS coil design that can deliver deep and spot size adjustable stimulation to deep brain regions.

**Methods:** In our design, we introduce a magnetic core at the center of a coil to help confine the magnetic field and prevent leakage. We further tilted the wire wrapping angle of the coil to break its ring symmetry and accomplish tunable focusing by adjusting the tilting angle.

**Results:** By comparing the electric field decay curves of five types of coils, our results concluded the proposed novel method to improve the coils’ depth-focality profile. Both theoretical calculations and experimental data collectively demonstrated that by using a larger tilting angle, we were able to accomplish a more tightly focused stimulation at any distance away from the coil.

**Conclusion:** Enlarging the tilting angle of the coil wire wrapping and applying magnetic core significantly improved the spatial resolution of the field without inducing considerable effect on field decay speed. Our novel TMS coil design plots a new curve in the depth-focality profile with better performance than the existing conventional coil designs in the tradeoff rule.

## I. Introduction

Transcranial magnetic stimulation (TMS) has been approved by the Food and Drug Administration (FDA) for treatment-resistant major depression^1^ and Obsessive-Compulsive Disorder^2^. Its therapeutic effects in other psychiatric and neurological disorders, including drug addiction, are emerging ^3,4^. From both clinical and basic neuroscience perspectives, there has been a strong demand to obtain stimulation tools that can reach deep brain regions with small size targeted stimulations. For example, decades of neuroimaging studies have identified malfunction of dorsal anterior cingulate cortex, insular and amygdala in a range of psychiatric disorders. These structures are 4 cm or more below the scalp. Unfortunately, with current technologies, the stimulation targets are largely limited to the cortical surface, or otherwise large brain areas are stimulated when a deep brain structure is targeted.

The output of a TMS coil can be treated as field emission from a finite size aperture and follows a specific depth-focality tradeoff rule. Deng et al. theoretically calculated the depth-focality profiles of 50 TMS coil designed for stimulating a head model with diameter of 17cm ^5^. Two groups of mainstream coils, circular and the figure-8, formed two depth-focality tradeoff curves, respectively. The study concluded that at shorter depth, which is smaller than 7cm, all figure-8 type of coils follows a better depth-focality tradeoff rule and it will be advantageous to use figure-8 coil. A number of studies have attempted to design TMS coils for enhanced penetration depth or improved focality. Rastogi modified conventional the figure-8coil to improve focality but that, on the other hand, significantly weakened the electric field strength generated in brain tissues ^6^. Crowther suggested a “Halo coil” design to improve the penetration depth of common circular coils ^7,8^, but this design sacrifices the coil focality. Luiz modeled multichannel coil arrays to improve the focality and penetration depth profile ^9^. However, this design involved complicated coil structures, so that it required higher performance of the coils’ cooling system. Alternative coil design strategy is needed to go beyond the depth-focality tradeoff limitation. Roth et al. have developed the H-coil for human deep brain stimulation, but the design is still limited by the depth-focality tradeoff with a relatively large focality ^5, 10, 11^.

We recently reported a coil design approach that employed a long magnetic core (15 cm); the current sources were distributed vertically around the core, as opposed to horizontally on the same plane commonly seen in most commercial designs ^12^. The magnetic core confines magnetic flux and reduce field leakage. The focality of the electric field was further improved by a winding strategy that breaks the symmetry of the B field distribution. The penetration depth of this coil, however, was limited to a few millimeters, which was demonstrated by in-vivo data ^12^. The goal of this study is to extend this novel strategy to design TMS coils that reach deeper targets in the brain while maintaining high focality for non-human primate and human brain stimulation. Theoretical studies were performed with Finite Element Modeling (FEM), confirmed experimentally by field mapping of prototype coils. The field strength and decay rate of multiple coil structures were also compared with our novel design. Our data reveals a novel approach for the development of next generation coils critical for focal TMS of deep brain structures.

## II. Theoretical Background

Mathematically, the depth-focality tradeoff can be described as field distribution produced by an aperture of fixed size. The field produced by an aperture, as shown in Figure 1 (blue area), at a specific distance (*r*) from the aperture source can be treated as a summation (or integration) of vector field generated from all the point sources at the aperture as shown in Equation (1). The integral of field superposition from all the point sources at the aperture can be simplified to a mathematic form similar to the Fourier transform for either far or near field cases ^13^ as shown in Equation (2).

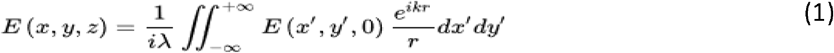

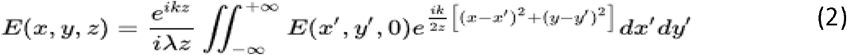

**Figure 1.**
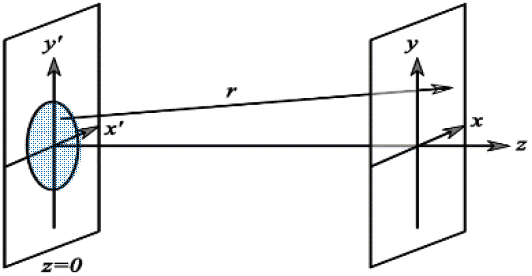
is a simplified illustration of the depth-focality tradeoff. The electric field produced by an aperture can be treated as a summation of the vector field generated from all the point sources at the aperture (Equation (1)), which can always be further simplified to Equation (2), similar to a Fourier transform.

Therefore, the aperture size (x) and the field distribution solid angle (k) follows x-k Fourier transform relationship, which means the larger the aperture, the smaller the field spread. One implication from Equation (2) is that by using large diameter coils, one can obtain slower decay of the emitted E field. Indeed, the theoretical study by Deng et al. identified an intersection of the two depth-focality curves arising from two general coil geometries: the figure-8 and circular. ^5^ The field generated by a figure-8 coil has a smaller equivalent aperture than that of a circular coil, since it is produced from the summation area of two ring coils. On the other hand, the circular coil induces ring shape electric field distribution, which can be large in regions in close proximity to the coil’s dimension. But the divergent angle can be smaller than that of a figure-8 coil. Based on the above theoretical analysis, we chose the circular coil geometry for deep brain stimulation.

## III. Results

### Modeling of field strength and decay rate for 5 basic TMS structures

Based on Equations (1) to (2), in order to enhance electric field focusing and field strength, we propose a coil design as follows in Figure 2(a): in contrast to conventional air-core coil design (A, B), we apply a magnetic core to the center of the coils (C, D), and we allow a tilt angle between the conductor and the magnetic core (E) to further improve electric field focality.

**Figure 2.**
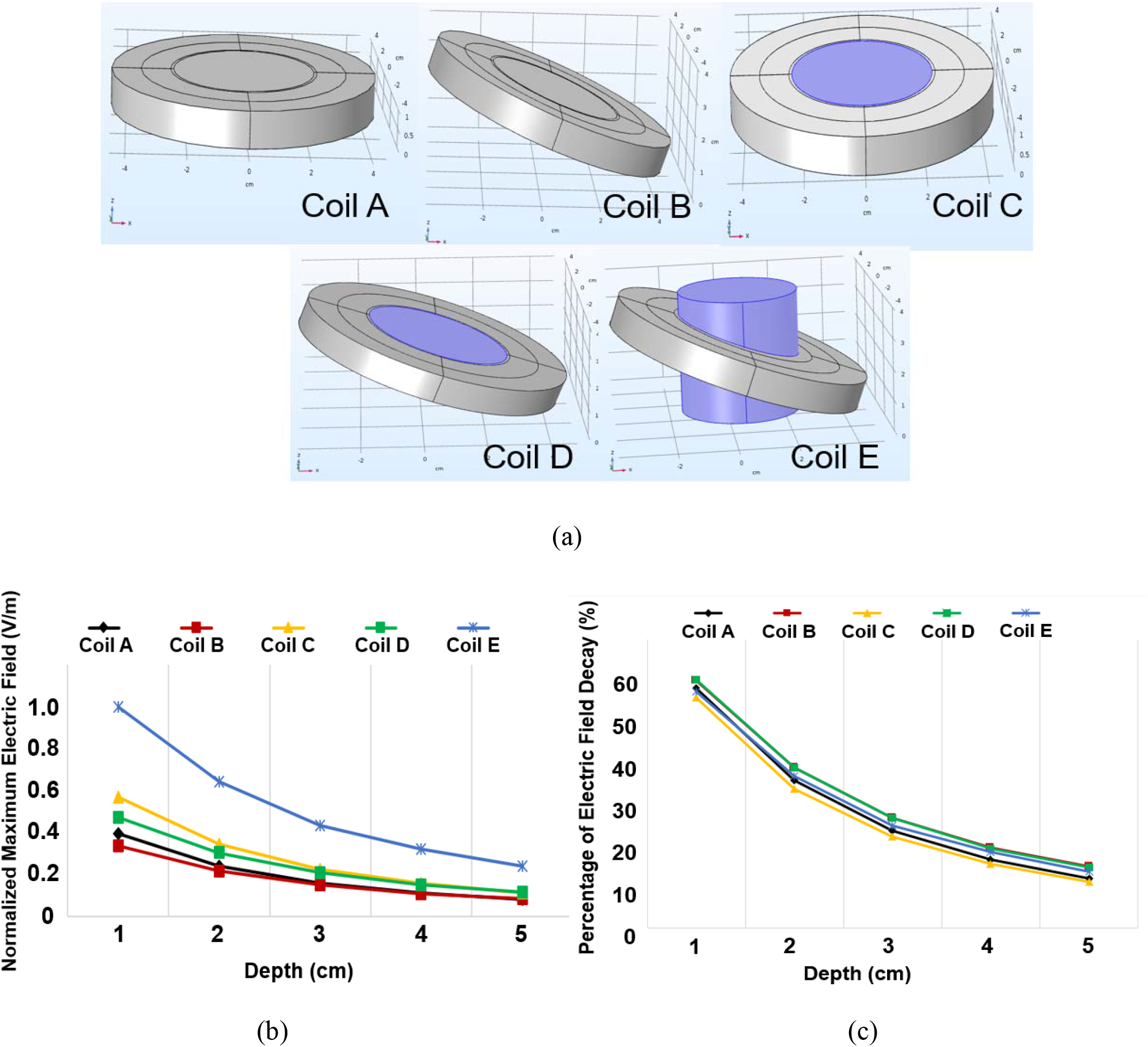
(a). Comparison of 5 models of TMS coils. Coil A: Air core human TMS coil, scanned planes are in parallel with the coil down-side surface; Coil B: Coil A tilt by 20 degrees along X axis, scanned planes have a 20 degree angle with the coil down-side surface; Coil C: Iron core human TMS coil, scanned planes are in parallel with the coil down-side surface; Coil D: Coil C tilt by 20 degrees along X axis, scanned planes have a 20 degree angle with the coil down-side surface; Coil E: Based on Coil D, but the magnetic core is not tilt with the coil turns, scanned planes are still in parallel with the core down-side surface (For all coils, the inner and outer diameters are 5cm and 9cm, respectively; thickness is 1cm.); (b). Comparison of field strength at multiple depth levels for Coils A to E; (c). Comparison of field decay rate at multiple depth levels for Coils A to E.

We modeled the field distribution of the above 5 coil designs using the COMSOL software package (finite element analysis software, COMSOL Inc). All coils shared the same dimensions: the inner and outer diameters were 5cm and 9cm respectively, and the coil thickness was 1cm. Each coil contained 8 turns of conductors. The current in each turn was set to the same amplitude with the frequency of 1MHz, since the rising edge of a typical TMS pulse is a few microseconds. The relative permeability of the magnetic core was set to 4000 as a constant and its conductivity was set to 0S/m to avoid the influence from the eddy currents in the core generated by the time-varying magnetic field. Induced electric field distributions generated by each coil were calculated at multiple depth levels (within the X-Y planes with a variable of the Z coordinate) and the decay rate along the Z direction was compared.

Figure 2(b) and (c) compares the normalized electric field strength and field decay rate at multiple depth for all 5 coil models. For air-core Coils (A, B), with a 20-degree tilting, Coil B delivers relatively lower field strength at each depth compared with Coil A (no tilting); but improves field decay rate along the Z direction by about 3%. With the help of the magnetic core, Coil C enhances field strength by 40-50% than Coil A (air core). As the depth gets deeper, this gain gradually reduces. In addition, the magnetic core accelerated field decay rate by 1% to 3% at each depth level along the Z. If the core was tilted together with the coil winding (Coil D), the decay rate would recover and be comparable with Coil B. But compared with Coil C, the electric field induced by Coil D is reduced by 20% and 25% at a depth of 1cm and 2cm, respectively. This effect gets weaker at deeper targets and eventually disappears at the depth of about 5cm.

Coil E features a tilt angle between the vertical core and the winding, and the core extends to the same height of the winding. This design substantially enhances field strength at all depths compared with Coils A-D, as shown in Figure 2(b). More specifically, compared with Coil C (the 2^nd^ best design), Coil D enhances the gain by a factor 1.8 at the depth of 1cm, increasing to 2.1 when the depth reaches 5 cm. Notably, all 5 designs share similar E field decay rate, as shown Figure 2(c). Thus, amongst the 5 designs, Coil E is most promising for deep brain stimulation. We next investigate the effects of other design parameters: coil length and tilt angle, on the strength and focality of the electric field in this design.

### The effect of coil length on electric field

We next modeled the coil length as a linear function of the number of single coils accumulated along the Z axis; the length of the magnetic core was extended accordingly. We kept the tilt angle at 20 degrees. As the number of accumulated coils increases from 1 to 7, the E field increases superlinearly. (Here one coil means one model of Coil A or Coil B in Figure 2(a), which contains 8 turns.) We quantified the gain to be G=α×N^2^, where α is a positive constant. Results are shown in Figure 3 (blue curve). For comparison, we also modeled the coil design without the magnetic core. The electric field tends plateau as the coil length extends beyond 3 cm (Figure 3, orange curve). This curve can be well-fitted with G=β×N^0.5^, where β is another positive constant.

**Figure 3.**
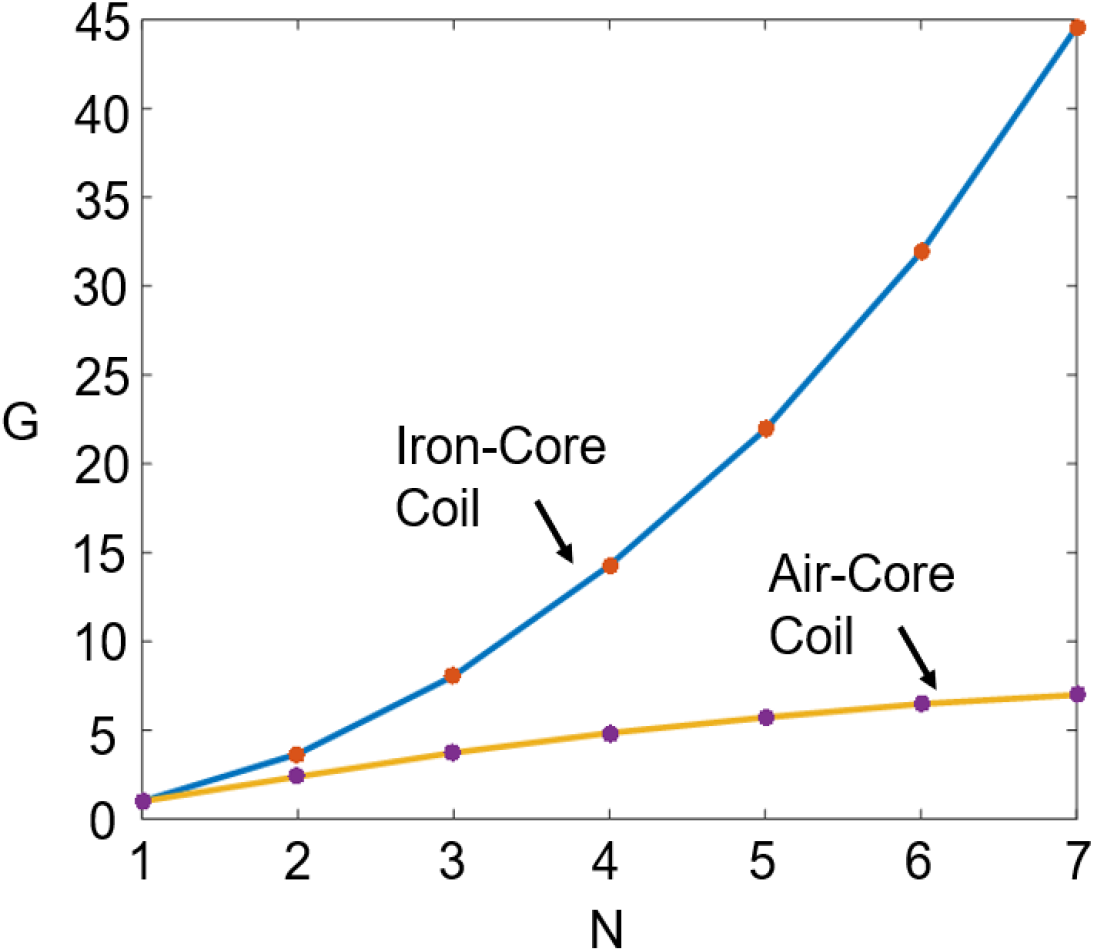
Gain effect of coil turns accumulation along the Z axis for both iron-core coil and air-core coil. N is the number of single coils as shown in Figure 1(a) (Coil A) accumulated along the Z axis and G is the gain of maximum induced electric field strength by multiple accumulated coils at the depth of 2cm compared with only one coil in the model.

### The effects of the tilt angle on TMS focality

The conventional circular TMS coil features the B field that is circularly symmetric along the Z; the resulting E field is zero in the center and peaks in a circular plane, leading to poor focality. The introduction of a tilt angle (Coil E in Figure 2) breaks the circular symmetry in the B field, the resulting E field peaks in a focal region rather than a circular plane. To further investigate how the tilt angle affects the focality, we extended Coil E with 8 coils accumulated along the Z axis to about 15cm in total length and adjusted the angle θ from 0 to 10 and then 20 degrees. As shown in Figure 4(a), COMSOL simulation results clearly suggest that a larger tilt angle of wire wrapping achieves better electric field focusing at all depths. It is also found that with a larger angle (typically over 30 degrees), a considerable electric field component along the Z axis would be detected. The electric field distribution is then no longer within a two-dimensional plane, which is in parallel with the X-Y plane.

**Figure 4.**
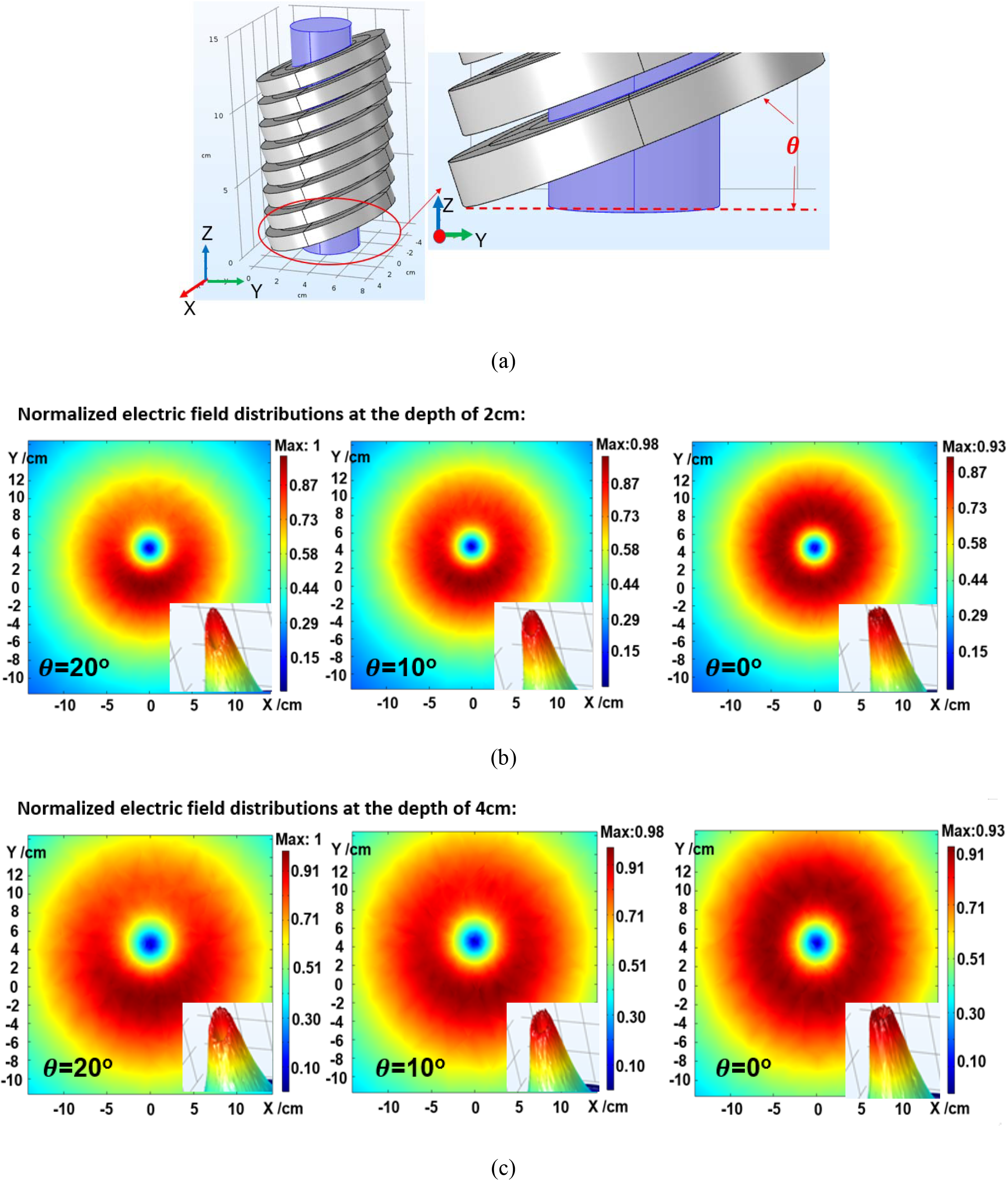
(a). COMSOL model of TMS coil with tilt wire wrapping; (b). Electric field distributions at the depth of 2cm for tilt angles of 20 degrees, 10 degrees and 0 degree (flat coil); (c). Electric field distributions at the depth of 4cm for tilt angles of 20 degrees, 10 degrees and 0 degree.

Figure 4 (b) and (c) compares field distributions at the depth of 2cm and 4cm of the three coil models in both two-dimensional and three-dimensional plots. The coil with 20-degree tilt angle forms a smaller area of focal spot (dark red area) than the 10-degree tilted coil. At the same depth, 20-degree coil also induces relatively stronger electric field intensity at the peak of its focal spot than the other two coils. The focusing effects continuously exist along the Z direction for a depth of more than 5cm. Similar to the iron cores inserted in the animal TMS coils ^12^, the iron cores in the human TMS coils also absorb magnetic flux in the coil and significantly reduced field-leaking. The electric field focal spot is guided by the core to a desired target aligned with coil dimension instead of to the direction of wire wrapping tilt, which follows the principle presented by Coil E in Figure 2(a).

### Verification of the electric field distributions in 3 coil prototypes

To verify the FEM simulations shown in Figure 4, we made 3 coil prototypes with tilt angles of 0, 10 and 20 degrees, respectively. They shared the same coil length, inner and outer diameters, which were 4.4cm, 3.8cm and 7.5cm respectively. Each coil was wrapped 20 turns of the litz wires, each turn contained a bundle of 135 piece of AWG30 wires. The TMS coil was driven by a customized driving circuit, in which an insulated gate bipolar transistor (IGBT) was used as the switch to control TMS pulses and a capacitor bank charged by a power supply was the main current source ^12^. The charging voltage of the capacitor bank was set to 100V and pulse duration was 250µs. The induced electric field was measured with a modified Rogowski E-field probe customized in our lab.^14,15^ The E-field was mapped at 2cm away for coil surface. Since the electric field components along the Z axis was small enough to be negligible, only the X- and Y-components of the E-field were measured. Considering the probe size, we mapped the E-field at a step size of 5mm within the X-Y plane. An area of 8cm×8cm was scanned for each coil.

Figure 5 shows heat maps of the E-field distributions. For the coil with a tilt angle of 20 degrees, the area with the field strength ≥ 80% of its peak value (E_peak_) is only 14.5 cm^2^; while for the other 2 coils, the values reach 22.5 cm^2^ (for 10-degree tilt) and 30.75cm^2^ (for 0 degree tilt), respectively. Thus, 2 cm away from coil surface, a tilt angle of 20 degrees improves the E-field focality by a factor of 2.6 than a flat TMS coil. The experimental results match well with COMSOL simulations. It is also found that a larger angle would slightly increase the coil’s inductance. For example, the inductances of the 3 scanned coils were measured to be 52.2uH, 51.21uH and 49.4uH for 20-degree, 10-degree and 0-degree wire wrapping, respectively.

**Figure 5.**
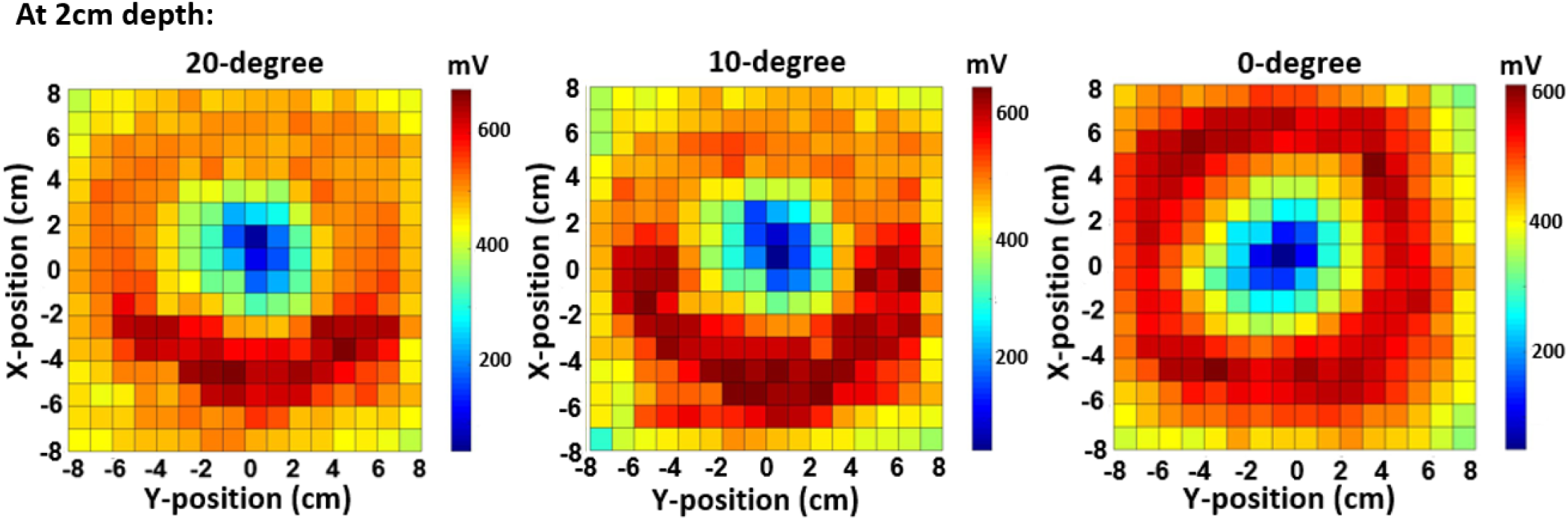
Induced electric field measurements using modified Rogowski coil probe for the 3 coils with 20 degrees, 10 degrees and 0 degree wire wrapping at the depth of 2cm.

## IV. Discussions

The combination of iron core and the tilt coil wrapping provides a significant innovation to the depth-focality profile of TMS coils. The tilt angle technique is a mild symmetry breaking method. It does not reduce the equivalent field emission aperture size or speed up the decay rate along the Z direction, but significantly distorts the ring shape electric field distribution, resulting in a much smaller focal spot. Application of an inserted magnetic core was able to replenish the loss of field created by the tilting without affecting field decay considerably. It was guiding the magnetic flux with stronger density to the core axis direction, which was not aligned with the tilt angle direction. With the limitation of the depth-focality tradeoff, ^5^ our novel TMS coil design provides the current best solution to improve this profile by dramatically shrinking the focality and simultaneously maintaining nearly the same decay rate of the induced electric field strength. As the stimulation target gets deeper, the circular type of coils present better performance than the figure-8 type coils in the depth-focality tradeoff profile. In the study of 50 coils’ electric field depth-focality tradeoff by Deng et al., the authors detected an intersection of the depth-focality curves for these two types of coils when the half-depth value reaches 3.5cm. ^5^ This intersection indicates a deeper stimulation depth beyond this point, and the profile curve of the circular coils will be below the figure-8 coils. Our novel design does not obviously change the decay rate of the conventional circular coils. On the other hand, significantly reduces the focal spot size, which provides the potential to reach deeper region when the coil diameter is enlarged.

The only limitation of the angle-tuned TMS coil design, to our knowledge, is the relatively larger inductance compared with conventional TMS coils. The current flowing inside the coil I(t) can be expressed as

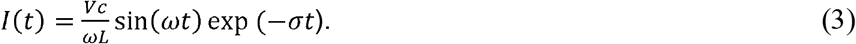

Here ω and σ are related to the charging capacitors (C), the inductance of the coil (*L*), and the resistance in the LC circuit. *V*_*c*_ is the voltage of charging capacitor. ^16^ The induced electric field *E(t)* can be expressed as 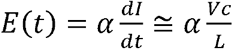. So, *E* is inversely proportional to *L.* The Magstim figure-8 human TMS coil has an inductance of about 17 μH (from the Magstim Rapid2 system manual). Our prototype human TMS coil has an inductance of 52 µH. This would require a stimulator of higher power to drive the coil. On the other hand, the use of magnetic core can drastically enhance the *B* field. This, in principle, reduces power requirement of the stimulator.

## V. Conclusion

A new angle-tuned TMS coil design is proposed and demonstrated. It can provide adjustable stimulation spot size by changing the wire wrapping angle, and at the same time keep the equivalent emission aperture the same. The long ferromagnetic core prevents magnetic leaking and superlinearly increases the field strength as a function of the coil length. It is also found that the tuning angle will not affect the coil’s depth dependent field-decay rate. As a result, the proposed coil design would theoretically present a much better performance in the depth-focality profile by Deng et al. than conventional TMS coils at deeper stimulation depth ^5^. This novel design provides a promising solution for future deep and focused human brain stimulations.

## Acknowledgement

The research is supported by the intramural research program of NIDA, NIH, NSF grant ECCS-1631820, NIH grants MH112180, MH108148, MH103222, and a Brain and Behavior Research Foundation grant.

